# CardioSeg: An interactive platform for integrated spatial transcriptomics data and nuclear morphological analysis of mouse heart tissue

**DOI:** 10.64898/2026.05.21.723734

**Authors:** Srijan Karthik Kancherla, Arne Olav Melleby, Jan Magnus Aronsen

## Abstract

**Motivation:** Spatial transcriptomics enables gene expression profiling within its spatial context in intact tissue sections. Existing workflows for segmentation, spatial annotation, and morphological analysis are often code-heavy and poorly integrated. This limits the joint analysis of spatial gene expression at a single-nucleus resolution, and corresponding nuclear morphology.

**Results:** We present CardioSeg, an integrated computational platform for nuclei-resolved spatial transcriptomic analysis combining multi-threshold segmentation, transcriptomic aggregation, cell-type annotation, and interactive morphometric querying within a unified graphical interface. CardioSeg achieved robust segmentation performance across heterogeneous imaging conditions, with union-based inference outperforming the individual parameter configurations. CardioSeg achieved 0.88 in accuracy and 0.85 in balanced accuracy against reference labels, while also resolving spatial heterogeneity not captured by spot-based approaches. Analyses of pressure-overloaded cardiac tissue indicated altered cell composition, nuclear morphology and gene expression in specific segments, indicating the potential of CardioSeg to couple disease-specific nuclear morphology with the associated transcriptomics.

**Availability and Implementation:** Source code is available at GitHub under the CC BY 4.0 license (https://github.com/SrijanKancherla/CardioSeg). A versioned release was archived in Zenodo (DOI: 10.5281/zenodo.20177171).

## Introduction

The era of single-nucleus sequencing combined with spatial transcriptomics (ST) has transformed the study of cellular biology in the heart and other organs and allows analyses of cell neighbourhoods and cell-cell interactions within intact tissue (Litvinukova et al. 2020; Kanemaru et al. 2023; Palmer et al. 2024). Until recently, the resolution of ST data has been lower than the single-cell size, requiring computational deconvolution and preventing direct mapping of transcriptional profiles to individual cells in histological sections.

Advances in high-resolution ST have overcome this limitation by enabling near single-cell or subcellular resolution, allowing the direct coupling of spatial signals to individual cells from corresponding histology sections. This allows for more precise identification of low-abundance cell types as well as mapping of gene expression and morphology at the level of individual nuclei. Existing pipelines for the analysis of ST data often require multiple tools and code-intensive workflows to perform segmentation, annotation, and downstream analysis. Widely used platforms such as CellProfiler (Carpenter et al. 2006; Dao et al. 2016) and QuPath (Bankhead et al. 2017) provide powerful capabilities but require substantial computational expertise and lack integrated visualization, limiting iterative exploration.

Emerging evidence indicates that nuclear morphology, such as area (Anversa et al. 1991; Gerdes and Capasso 1995; Mollova et al. 2013), circularity (McCain et al. 2013; Otsuji et al. 2010), and eccentricity (Fatkin et al. 1999; Muchir et al. 2007), is an indicator of cardiac pathology. However, the lack of tools for nuclear segmentation of myocardial tissue hinders the integration of morphological features with spatial transcriptomic profiles.

Here, we present CardioSeg, a Python-based graphical user interface (GUI) designed for nuclei-level spatial analysis of cardiac tissue. By unifying segmentation, annotation, and interactive analysis on a single platform, CardioSeg provides an accessible and scalable solution for high-resolution spatial biology, enabling both hypothesis-driven and exploratory investigation of cardiac tissue organization. The platform incorporates a multi-threshold segmentation strategy for robust segmentation across heterogeneous imaging conditions, along with a marker-based annotation algorithm for assigning cell identities. Additionally, CardioSeg introduces a query language that enables real-time filtering, visualization, and statistical interrogation of morphological and transcriptomic data for users with limited programming experience.

## Materials and methods

### 1. Overview of CardioSeg

CardioSeg couples high-resolution microscopy images of haematoxylin and eosin (H&E)-stained myocardial samples with corresponding ST data as input, performs nuclei segmentation, and assigns cell-type annotations to nuclei across input images (Figure 1). Next, CardioSeg provides an easy-to-use GUI that allows the user to explore regions-of-interest (ROIs), such as vessels, the septum, free wall, endocardium, or related anatomical regions. Interactive querying and outputs include annotated cell type, gene expression, and nucleus morphology in individual nuclei. The software version used in this study was archived as a stable release on Zenodo.

**Figure 1:**
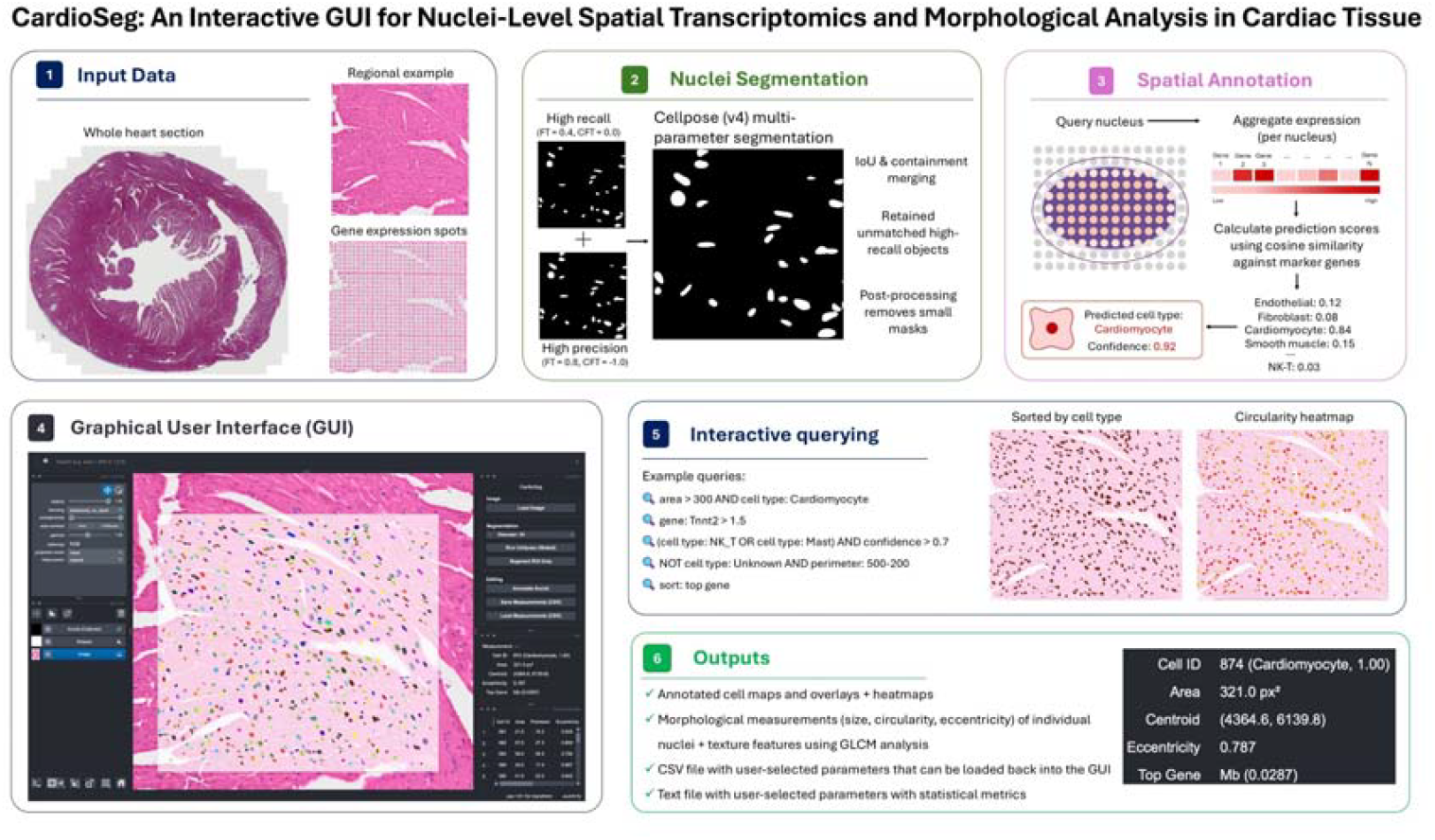
Visual summary of CardioSeg. See text for details. Colour coding in the image in part 4 is the default RGB spectrum to differentiate between nuclei in the selected region-of-interest.

### 2. Preparation of datasets: histology slides and spatial transcriptomics data

#### 2.1 Preclinical model, surgery and phenotyping

Sham or O-ring aortic banding (ORAB) surgery (Melleby et al. 2018) was performed to induce cardiac remodelling in ∼25g male C57BL/6JRj mice (Janvier, France). 21 days after surgery, echocardiography was performed using a Vevo 3100 system (FUJIFILM VisualSonics, Canada), and the left ventricles (LVs) were harvested under deep surgical anaesthesia (Skogestad et al. 2023). LV samples were washed in phosphate-buffered saline (PBS) and sectioned at the midventricular level along the short axis. Samples were fixed in 10% formalin for 24 hours, embedded in paraffin, and sectioned at 4 µm. All animal procedures were conducted in accordance with the Guide for the Care and Use of Laboratory Animals (NIH Publication No. 85-23, revised 2011, US) and were approved by the Norwegian National Animal Research Committee (FDU 30837).

#### 2.2 Preparation of histology slides and spatial transcriptomics

H&E staining and imaging were performed according to the protocol described in the Visium HD FFPE Tissue Preparation Handbook (CG000684). Following staining and imaging, tissue sections were processed and sequenced in accordance with the Visium HD Spatial Gene Expression Reagent Kit User Guide (CG000685) at the Genomics Core Facility of Oslo University Hospital.

### 3. Segmentation

Nuclei segmentation was performed using Cellpose v4 on high-resolution microscopy images (Stringer et al. 2021). Two Cellpose inference configurations were used to balance sensitivity and precision. A high-recall setting (flow threshold 0.4, cell probability 0.0) was used to capture faint or partially visible nuclei, and a high-precision setting (flow threshold 0.8, cell probability −1.0) was used to reduce the number of false positives. Both configurations produced instance masks that were merged using intersection over union (IoU) and containment-based rules. Overlapping detections are consolidated, nested masks are removed, and unmatched detections from the high-recall configuration are retained.

Post-processing removed small artifacts (radius < 5 pixels) and relabelled the instances.

For each segmented nucleus, morphological, textural, and intensity-based features were computed from the instance mask and the corresponding image, including area, perimeter, eccentricity, texture homogeneity, solidity, and circularity.

#### 3.1 Segmentation evaluation

The segmentation performance of CardioSeg was evaluated using five NuInsSeg images of myocardial samples (146 nuclei), and the masks predicted by CardioSeg were compared with NuInsSeg images with ground-truth annotations (Mahbod et al. 2024). Pixel overlap was quantified using IoU and Dice coefficients, and instance-level performance was assessed using F1-score, where the predicted and ground-truth objects were matched with an IoU threshold of 0.5. To assess the robustness to imaging resolution, segmentation was repeated on downsampled (512×512) images. All metrics were computed for each image and averaged across the datasets.

### 4. Cell Annotation

#### 4.1 Expression aggregation

ST data are represented as a gene expression matrix *x* ∈ ℝ ^*N*×*G*^, where *N* denotes the number of spatial spots, and *G* denotes the number of genes. Each spot *i* is associated with spatial coordinates *s*_*i*_ = (*x*_*i*_,*y*_*i*_). For each nucleus polygon *P*, spots were assigned using a KD-tree-based centroid search, followed by point-in-polygon filtering to obtain the set *s*_*P*_ spots within *P*. The aggregate expression profile for each nucleus was computed as:

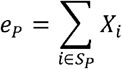

If no spots were contained within the polygon, the expression was estimated using inverse distance weighting over the *k*-nearest neighbor spots.

##### 4.1.1 Preprocessing and normalisation

Mitochondrial genes were excluded prior to analysis, and all expression vectors were normalized per cell.

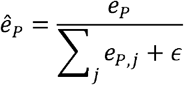

#### 4.2 Cell type assignment

Reference profiles *r*_*k*_ were derived from the Azimuth heart atlas (HuBMAP Consortium 2021), that integrates from scRNA-seq and snRNA-seq datasets from across multiple studies (Hocker et al. 2021; Litvinukova et al. 2020; Koenig et al. 2022) and mapped to mouse orthologs. The similarity between each nucleus and the reference profiles was computed using cosine similarity:

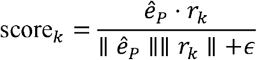

Cell identity was assigned as:

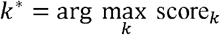

Confidence was defined as the separation between the top two cosine similarities.

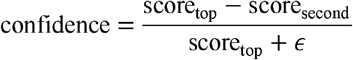

Cells with maximum similarity below a predefined threshold (0.2) were labelled as “*Unknown”*.

#### 4.3 Annotation evaluation

Annotation performance was evaluated using a publicly available mouse heart single-cell scRNA-seq dataset from 10x Genomics (2022). The predicted cell types were compared with ground-truth using accuracy, balanced accuracy, macro-averaged F1-score, and Cohen’s kappa. Dimensionality analysis was performed using Uniform Manifold Approximation and Projections (UMAPs). A marker-only baseline classifier was used to compare the transcriptomic annotations with the CardioSeg annotation engine. Both methods use cosine similarity to curated cardiac cell marker gene profiles; however, CardioSeg additionally incorporates spatial context through neighboring spatial transcriptomic spots.

### 5. Query Language for Interactive Spatial Cell Analysis

A domain-specific query language was used for nuclear morphology, gene expression, and annotation filtering. Each cell is represented as a structured object with an abridged dictionary as follows:

~~~
*Cell* {
   *id : int*
   *area : float*
   *centroid : (x,y*)
        …
   *eccentricity : float*
   *cell_type : string*
   *gene_name : float*
 }
~~~

Query syntax consists of logical and numerical expressions over cell attributes with supported operators, including

- Logical: *AND, OR, NOT*
- Comparison: >, <, >=, <=, =
- Range queries: : *feature:a* − *b*
- Categorical filters: *cell_type* : *Cardiomyocyte*
- Gene filters:
  - Presence: *gene* : *Tnnt2*
  - Quantitative: *gene* : *Tnnt2* > 0.3

Logical expressions can be combined using AND, OR, NOT, e.g.:

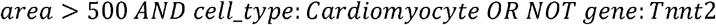

The queries are parsed into set operations using cell identifiers. Each condition is evaluated independently, and the results are combined using intersection, union, and complement operations. This approach enables an efficient evaluation across large datasets while maintaining consistency.

The results are mapped directly to visualizations using highlighting and feature-based overlays. The system also supports summary statistics for selected populations, including mean, median, and distributional analyses, with the optional export of results for downstream processing. To evaluate the nuclei in the remodelled myocardium, we used a herein proposed stress score that was calculated as follows:

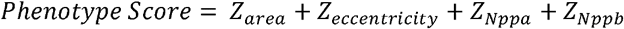

### 6. Statistics

All average numbers are reported as average value ± standard deviation.

## Results

### Evaluation of nuclei segmentation performance of CardioSeg

First, we evaluated the segmentation performance of CardioSeg on five annotated slides of myocardial samples from the NuInsSeg dataset, including 146 nuclei (Mahbod et al. 2024). The default Cellpose pipeline achieved consistent segmentation quality, and the individual parameter settings demonstrated a precision-recall trade-off. IoU ranged from 0.54–0.59 and Dice 0.70–0.74, with no single configuration consistently outperforming the others (Table 1).

**Table 1:** Segmentation performance across configurations. Cellpose v4 with default settings or flow and cell probability (CP) as indicated. IoU; Intersection over union.

| Method | IoU | Dice | Precision | Recall | F1 |
| --- | --- | --- | --- | --- | --- |
| Cellpose (default) | $0.560 \pm 0.050$ | $0.723 \pm 0.052$ | $0.664 \pm 0.108$ | $0.589 \pm 0.110$ | $0.630 \pm 0.105$ |
| Flow 0.4 / CP -1.0 | 0.555 | 0.711 | 0.650 | 0.536 | 0.587 |
| Flow 0.4 / CP 0.0 | 0.540 | 0.700 | 0.723 | 0.602 | 0.656 |
| Flow 0.8 / CP -1.0 | 0.593 | 0.744 | 0.611 | 0.622 | 0.617 |
| Flow 0.8 / CP 0.0 | 0.552 | 0.710 | 0.662 | 0.608 | 0.633 |
| **Union** | **$0.593 \pm 0.039$** | **$0.744 \pm 0.030$** | **$0.611 \pm 0.124$** | **$0.622 \pm 0.118$** | **$0.616 \pm 0.115$** |

The proposed multi-threshold union strategy combining complementary high-recall (flow threshold 0.4, cell probability 0.0) and high-precision (flow threshold 0.8, cell probability −1.0) Cellpose settings, achieved the most consistent performance (IoU 0.593 ± 0.039, Dice 0.744 ± 0.030) by reducing inter-image variability. To quantify its specific contribution, we compared the union output with the best-performing individual configuration using pixel-level difference analysis.

The union strategy primarily improved recall, recovering additional true-positive regions (4724 ± 439 gained pixels), while preserving existing detections (9591 ± 2875 stable pixels) (Figure 2). Figure 3 presents a qualitative comparison of the ground-truth and union predictions.

**Figure 2:**
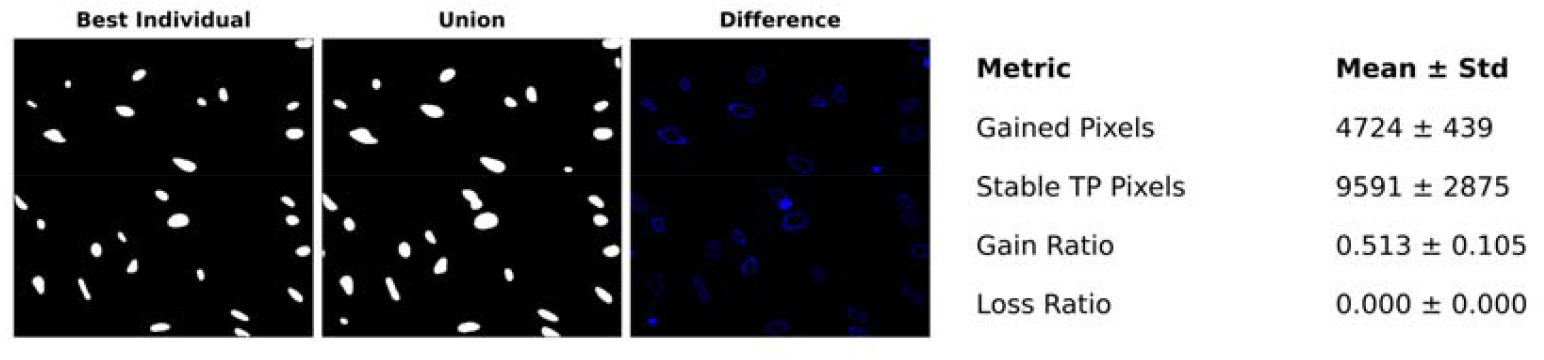
Comparison of the best individual method and the union method with a difference error map. Table shows pixel-level gain and loss analysis of union strategy.

**Figure 3:**
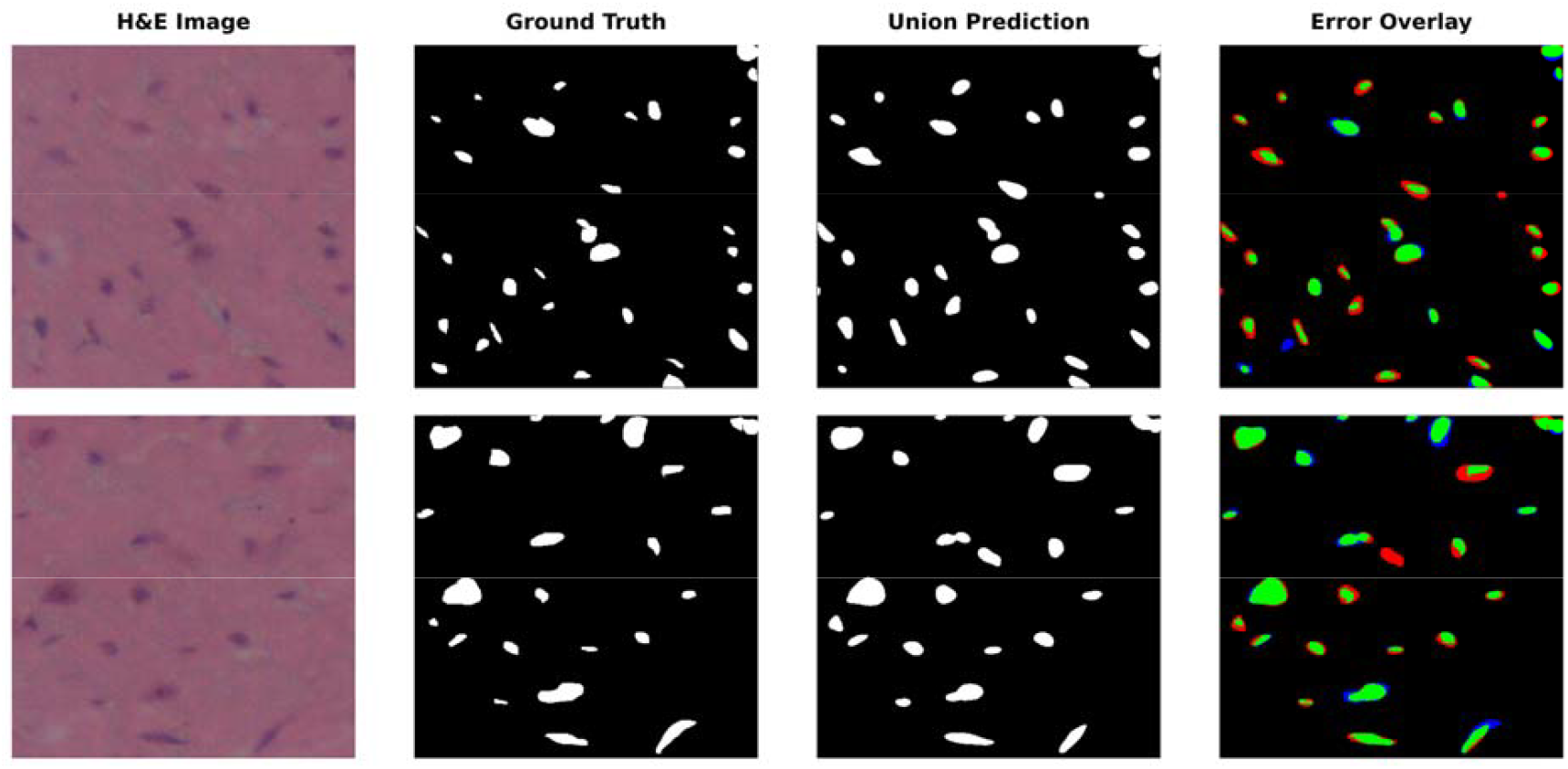
Two examples (upper and lower row) comparisons of ground-truth and union prediction masks with a coloured error overlay map using CardioSeg on NuInsSeg annotated slides of myocardial samples. Green, true positive; red, false positive; blue, false negative.

To evaluate robustness to image resolution, segmentation was performed on both high-resolution and downsampled (512×512) images across 909 nuclei. The performance decreased under downsampling (IoU 0.50 ± 0.19, Dice 0.65 ± 0.16), with increased false positives and missed detections (Figure 4).

**Figure 4:**
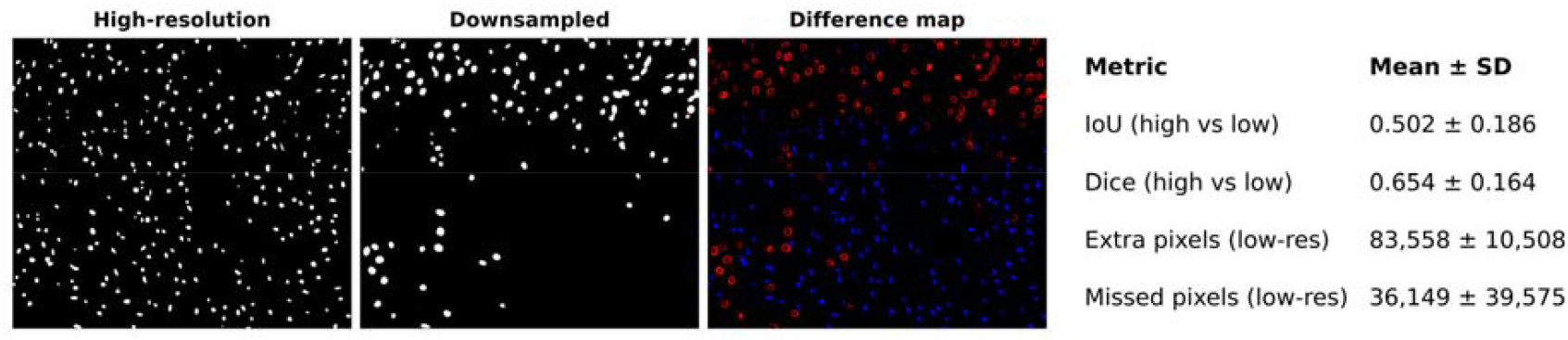
Effect of downsampling resolution of NuInsSeg annotated slides of myocardial samples on segmentation performance by CardioSeg.

Together, these results demonstrate that a simple multi-threshold ensemble strategy implemented in CardioSeg improves segmentation robustness, and that the remaining errors are primarily driven by boundary ambiguity and high nuclear density, exacerbated by reduced image resolution.

### Evaluation of cell type annotation by CardioSeg

Cell type annotation performance by CardioSeg was evaluated using a publicly available mouse heart single-cell RNA-seq dataset (10x Genomics, 2022), comprising 3282 cells. The proposed framework achieved an accuracy of 0.88, a balanced accuracy of 0.85, and Cohen’s kappa of 0.83 (Table 2). The macro F1-score was lower (0.48), primarily driven by the absence or limited representation of several low-abundance cell populations in the Visium spatial reference dataset, including lymphatic endothelial, mast, mesothelial, neuronal, and smooth muscle cells.

**Table 2:** Summary of cell type annotation performance by CardioSeg, evaluated on a mouse heart single-cell RNA-seq dataset with ground-truth labels from 10X genomics.

| **Metric** | **Score** |
| --- | --- |
| Accuracy | 0.88 |
| Balanced Accuracy | 0.85 |
| Cohen’s Kappa | 0.83 |
| Macro F1 | 0.48 |

A confusion matrix comparing cell type annotation by CardioSeg with 10X segmentation as the ground-truth is shown in Figure 5A. Marker gene heatmaps confirmed biologically coherent clustering and expected enrichment of canonical markers (Figure 5B). Compared to spot-based approaches, nuclei-level segmentation enables higher-resolution spatial annotation.

**Figure 5:**
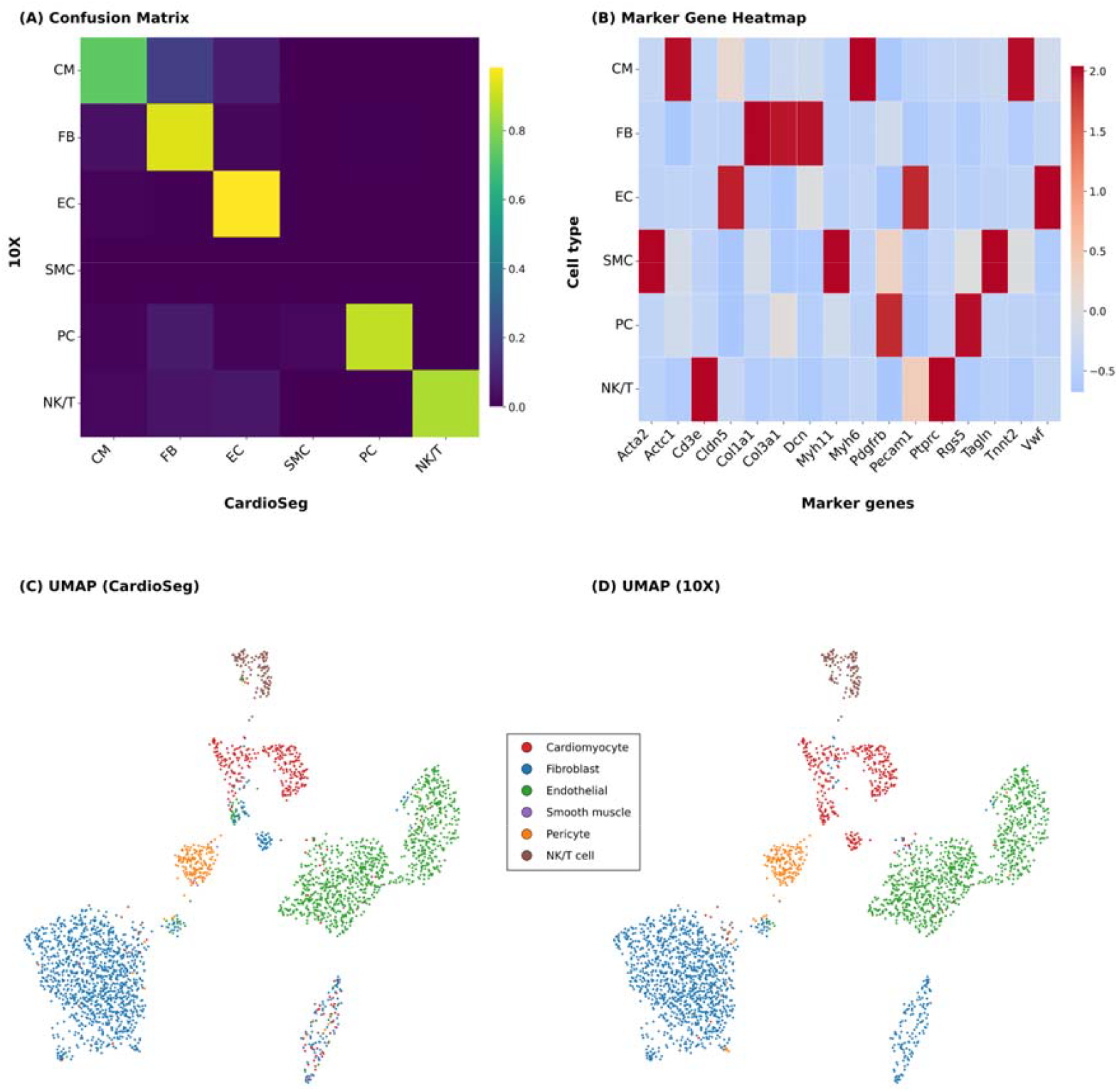
UMAP embedding, confusion matrix, and marker gene expression across predicted cell types. (A) Confusion matrix comparing CardioSeg prediction with 10X annotation from a single-nucleus dataset from mouse hearts as ground-truth. (B) Marker heatmap of genes in CardioSeg predicted cell types. (C, D) UMAPs of predicted cell types from CardioSeg (C) and 10X genomics annotated (D) cell types from the same dataset. CM = Cardiomyocyte, FB = fibroblast, EC = endothelial cell, SMC = smooth muscle cell, PC = pericyte, NK/T = natural killer T-cell

Major cardiac populations were consistently identified using CardioSeg. UMAP embedding revealed clustering into major transcriptional groups with partial overlap among finer subtypes (Figure 5C/D). Misclassifications primarily occurred between the immune and stromal populations, consistent with overlapping marker expression.

To assess the contribution of spatial aggregation, the method was compared to a marker-only baseline that assigns cell types without incorporating spatial context (Table 3). The marker-only baseline achieved a higher macro F1, but lower balanced accuracy and agreement. The spatial expression engine improved the balanced accuracy (0.849) and kappa (0.826), while increasing the assignment coverage (coverage: 3282 vs. 3039 nuclei).

**Table 3:** Comparison of marker-only and CardioSeg’s spatial expression engine annotation.

| Method | Macro<br>F1 | Balanced<br>Accuracy | Cohen's<br>Kappa | Coverage (n<br>samples) |
| --- | --- | --- | --- | --- |
| Marker-only baseline | 0.523 | 0.835 | 0.783 | 3039 |
| Spatial expression<br>engine | 0.475 | 0.849 | 0.826 | 3282 |

Together, these results demonstrate that the proposed annotation framework reliably captures major cardiac cell populations while revealing additional heterogeneity at nuclear-level resolution. The remaining ambiguities are primarily associated with transcriptionally overlapping cell types rather than with systematic limitation of the method.

### Visualisation of nuclei morphology and gene expression by CardioSeg

CardioSeg provides integrated visualization of segmentation, morphology, and gene expression within a unified interface. Spatial maps of nuclear morphology (area, perimeter, eccentricity, texture homogeneity, solidity, and circularity) and gene expression (e.g., *Tnnt2*) can be visualized directly on tissue images (Figure 6). Cell type annotations are displayed, along with their morphological and transcriptional features. This enables direct multimodal spatial exploration, without the use of external tools.

**Figure 6:**
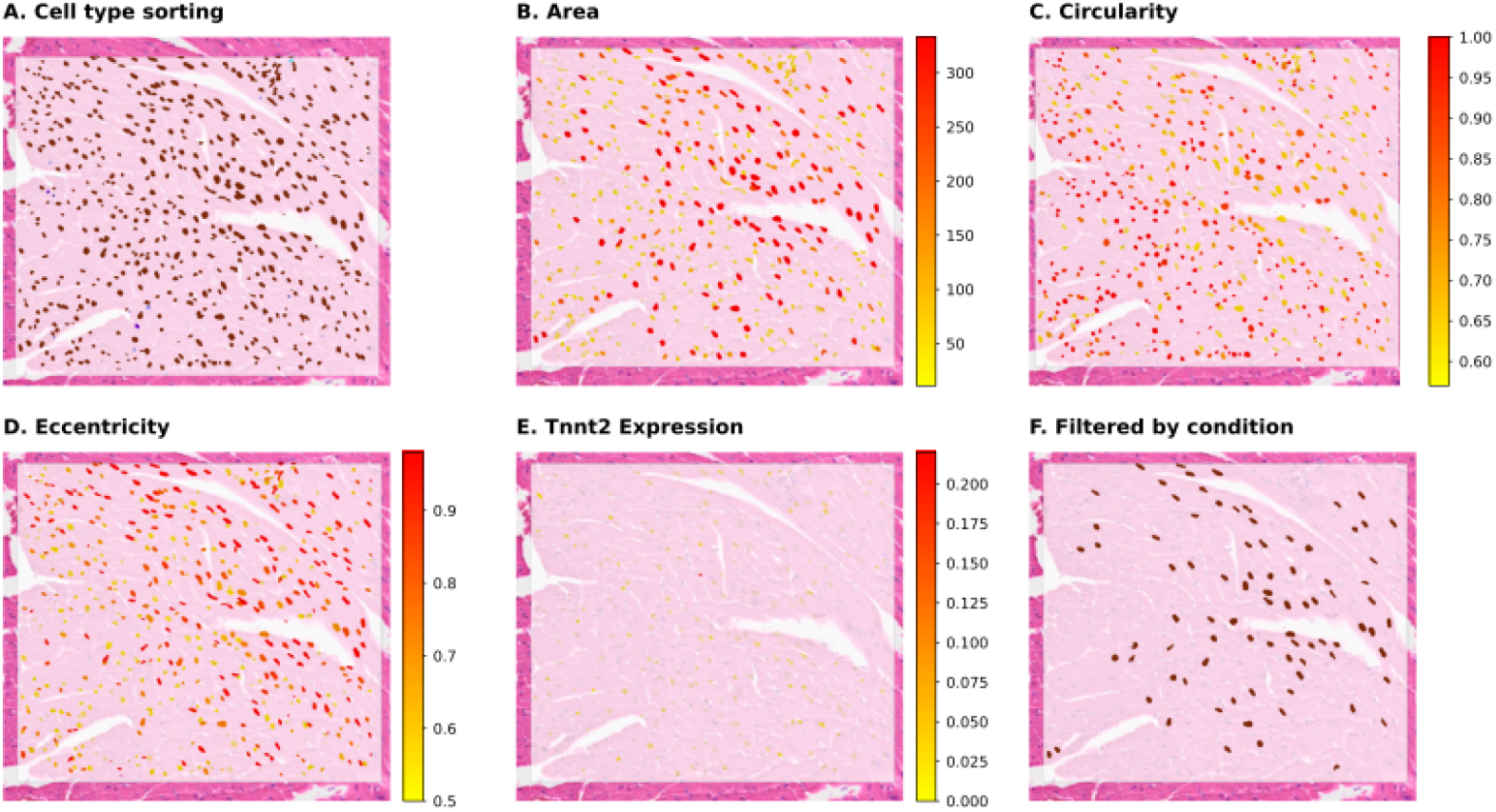
Visualization output showing segmentation and extracted nuclear morphological features and gene expression by CardioSeg. (A) Cell type sorting (red, cardiomyocytes; cyan, lymphatic endothelial cells; blue, fibroblasts). (B-D) Examples on visualization of area, circularity and eccentricity of nuclei in a mouse myocardial sample. (F) Filtered cells by “NOT cell type: Unknown AND perimeter: 50-200”.

**Figure 7:**
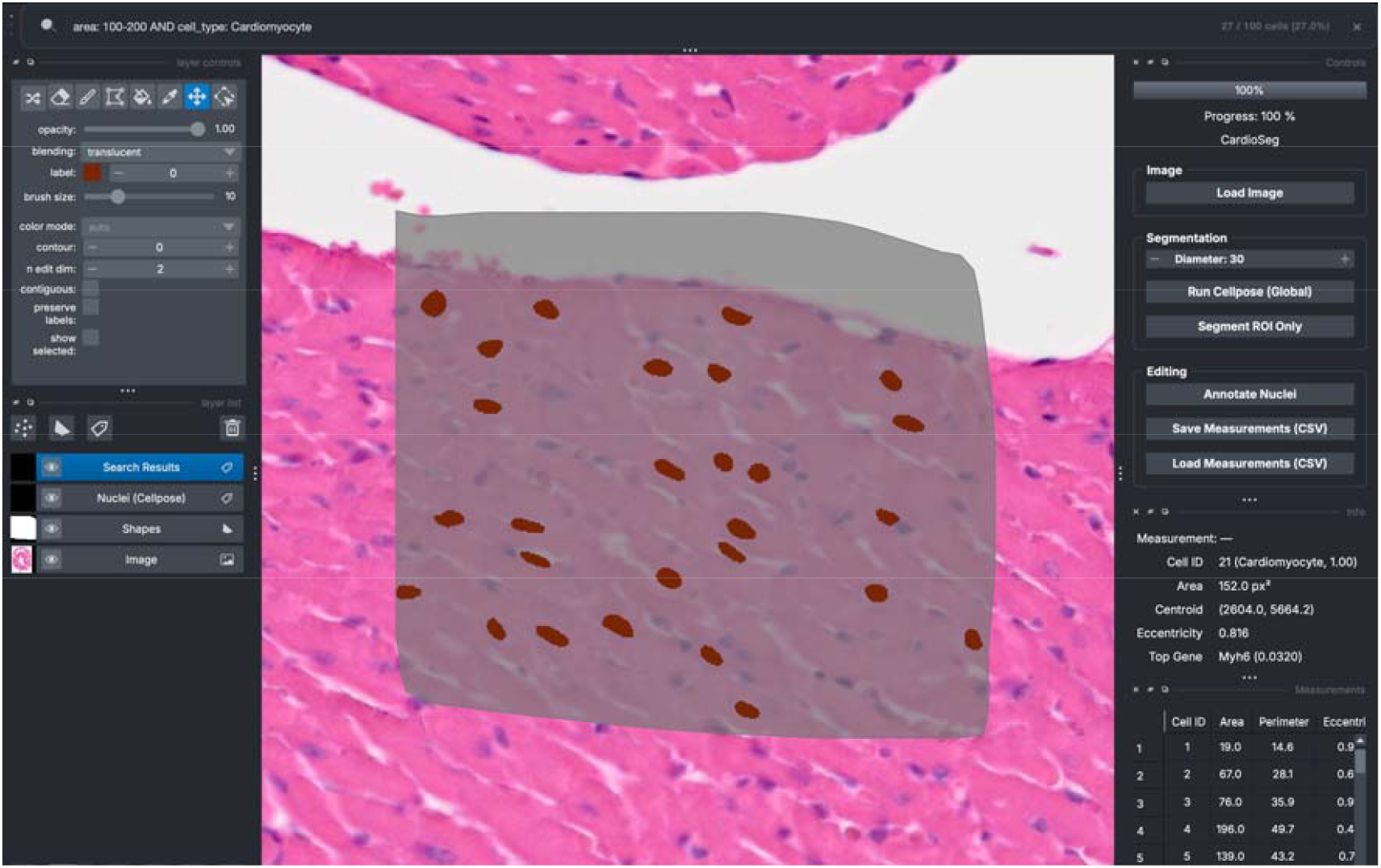
Example of interactive visualization and query-based filtering in the CardioSeg graphical user interface (GUI). The interface displays nuclei masks post-segmentation, cell type annotation, and morphological and transcriptomic directly shown on the side menu. The example shown illustrates filtered visualization following a compound query applied through the search bar.

CardioSeg enables the interactive filtering of nuclei-level features using a domain-specific query language. Users can filter by morphology, gene expression, and cell type in real time (example shown in Figure 6F). To evaluate usability, we compared the query system to existing spatial analysis platforms.

These results demonstrate that integrating segmentation, annotation, and querying enables the efficient spatial exploration of single-cell data (Table 4). By reducing the separation between data processing and visualization, the system supports efficient identification of spatial patterns and accelerates hypothesis generation.

**Table 4:** Comparison of query workflows across spatial analysis platforms.

| Feature | CardioSeg | CellProfiler | QuPath |
| --- | --- | --- | --- |
| Query interface | Domain-specific inline query | SQL-like filtering | Script-based (Groovy/Java) |
| Example query | area > 200 AND<br>gene: Tnnt2 | SELECT * FROM<br>per_object WHERE<br>Nuclei_Area > 200 | getObjects(...) |
| Execution model | Direct filtering with immediate visual update | Table generation followed by manual linking | Script execution with separate selection step |
| Visualization coupling | Integrated (automatic highlighting in viewer) | Indirect (requires mapping from table to objects) | Manual selection and visualization |
| Gene + morphology queries | Unified | Limited | Not native |
| Iterative exploration | Real-time | Multi-step | Multi-step |

### Integrated spatial profiling by CardioSeg

To demonstrate the capability of CardioSeg for integrated spatial analysis, we analysed left ventricular tissue from pressure overloaded mice subjected to ORAB or sham surgery 21 days post-operation. Region-resolved quantification of morphological and transcriptomic features was performed across the septum, hinge posterior (HP), and arterial compartments. Figure 8 summarizes SHAM and ORAB comparisons for selected genes and morphological features and reveals region-specific alterations in gene expression and nuclear morphology associated with pressure overload.

**Figure 8:**
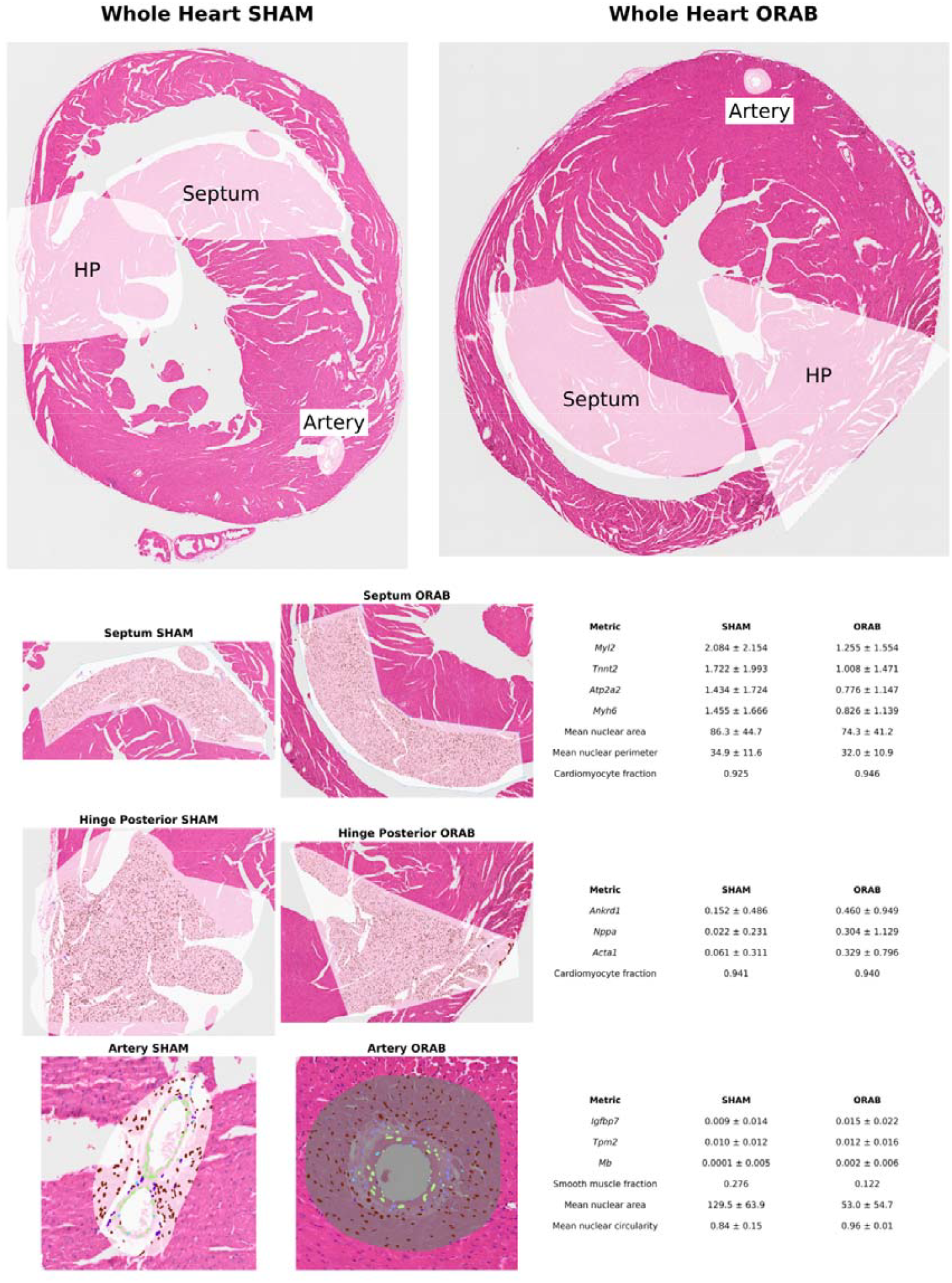
Integrated spatial transcriptomic and nuclear morphometric profiling of left ventricular samples 21 days after ORAB or sham surgery. Segmentation images are shown alongside quantitative summaries of selected gene expression and nuclear morphometric features across the septum, hinge posterior (HP), and arterial compartments. Gene expression analyses in the septum and HP were restricted to cardiomyocytes, whereas all arterial metrics were restricted to smooth muscle cells. Septal gene expression values are shown as raw measurements, whereas HP and arterial values are normalized. Nuclear morphometric analyses in the septum and HP included all segmented cells. Total nuclei analysed were as follows: HP SHAM, n = 5924; HP ORAB, n = 5361; Septum SHAM, n = 4909; Septum ORAB, n = 7798; Artery SHAM, n = 246; Artery ORAB, n = 312. The figure demonstrates CardioSeg’s integrated nucleus-resolved framework for simultaneous visualization of cell-type annotations, gene expression, and nuclear morphometric features within intact tissue architecture.

## Discussion

This article presents CardioSeg, an integrated framework for nuclei segmentation, spatial gene expression mapping, and interactive querying within a unified graphical interface. Validation metrics demonstrate reliable performance across segmentation and annotation while enabling integrated spatial exploration in a single system. This approach bridges the gap between image-based analysis and the spatial transcriptomic workflow.

Segmentation by CardioSeg is based on two Cellpose inference configurations (Stringer et al. 2021) combined using a multi-threshold union strategy, that is, a high-recall and a high-precision configuration to capture faint or partially visible nuclei and reduce false positives. Segmentation performance by CardioSeg showed consistent agreement with ground-truth annotations across myocardial samples in NuInsSeg (Mahbod et al. 2024), with improvements over the baseline Cellpose. These results should be interpreted in the context of the dataset’s resolution, which may underestimate the performance of the method. As we showed that lowering the microscopy resolution significantly increased false positives and missed detections, the use of CardioSeg should be based on high-resolution images.

Evaluation of the annotation performance of CardioSeg showed strong agreement with ground-truth labels, driven by the robust identification of major cardiac populations, including cardiomyocytes, endothelial cells, fibroblasts, pericytes, myeloid cells, and NK/T cells. This is reflected in the high global performance metrics; however, macro-averaged F1 is lower (0.48), consistent with substantial class imbalance and increased difficulty in resolving closely related or rare cell states. This performance is further explained by the structured misclassification among transcriptionally similar populations.

Compared to spot-based spatial transcriptomic approaches, CardioSeg enables nuclei-level resolution, allowing the detection of intra-cardiac heterogeneity and identification of additional endothelial, fibroblast, and immune-associated populations. Marker gene heat maps and embeddings confirmed that the predicted populations remained biologically coherent.

The integration of a query language within the GUI enables the real-time filtering, visualization, and statistical summarization of spatially resolved single-cell data. This is expected to facilitate the analyses of transcriptomic data from individual cells within their spatial context, such as cell niches and neighbourhoods, which have only recently been reported in the myocardium, despite their assumed fundamental role in cardiac remodelling (Kanemaru et al. 2023).

Nuclear morphology has been reported to be altered in various cardiac diseases, including increased cardiac hypertrophy (Anversa et al. 1991; Gerdes and Capasso 1995) and altered eccentricity with reduced elongation after increased load, and in heart failure. Additionally, irregularities in the nucleus may be indicators of nuclear membrane injury. CardioSeg enables direct comparison of nuclear morphology and spatial gene expression, which we believe will facilitate further studies on the coupling between cardiac disease, nuclear morphology, and transcriptomic responses. CardioSeg identified altered nuclear morphology following pressure overload that varied in different regions of the heart, highlighting the potential of CardioSeg to facilitate novel insights in cardiac remodelling by coupling nuclear morphology with spatial context and associated transcriptomes. A limitation of this approach is that nuclear size and morphology depend on the orientation of tissue sectioning relative to the axis of the elongated cardiomyocyte nuclei. For example, longitudinal and transverse muscle fibers contain nuclei with different orientations, which should be considered when interpreting nuclear morphology measurements from microscopy sections.

Here, CardioSeg analysed nuclei in anatomically defined compartments (septum, hinge posterior, and artery), allowing direct side-by-side comparison of structural and molecular readouts within the same interactive environment, sorting nuclei by cell type in samples from sham and ORAB-operated mice. This integrated layout enables visual interrogation of how nuclear structure and transcriptional programs can be analysed in various spatial regions and compared between experimental conditions. Importantly, all modalities are linked at a single-nucleus resolution, allowing users to dynamically connect segmentation output, cell-type assignment, and quantitative feature extraction within a unified interface. Together, this figure highlights the central design principle of CardioSeg: unifying segmentation, spatial annotation, and morphometric analysis in a single visual and computational framework for interactive exploration of cardiac spatial transcriptomic data.

Overall, CardioSeg demonstrated that combining segmentation, annotation, and querying within a single platform enables integrated analysis of cardiac tissue organization. CardioSeg shifts analysis from scripting-based workflows to an interactive environment, designed to be an easy-to-use platform accessible to users with limited programming experience. Future work will extend this framework to whole-cell segmentation, improve reference atlases for rare cell types, and integrate higher-resolution spatial transcriptomic data.

## Acknowledgments

The authors are thankful for the expert help provided by the Genomics Core Facility at Oslo University Hospital and the Department of Comparative Medicine at the University of Oslo.

## Author contributions

S. K. K. developed the theoretical frameworks and code, performed computations, and wrote the first draft of the manuscript. A. O. M. and J. M. A. conducted preclinical experiments and secured funding. J. M. A. supervised the study and revised the manuscript accordingly.

## Data access

The data underlying this study were derived from sources in the public domain: the NuInsSeg dataset, available on <u>NuInsSeg Kaggle page</u>, and the 5k Adult Mouse Heart Nuclei Isolated with Chromium Nuclei Isolation Kit dataset, available on <u>10x Genomics dataset page</u>.

## References

Anversa P, Olivetti G, Capasso JM. Cellular basis of ventricular remodeling after myocardial infarction. Am J Cardiol 1991;68:7D–16D.

Bankhead P, Loughrey MB, Fernández JA, et al. QuPath: Open source software for digital pathology image analysis. Sci Rep 2017;7:16878.

Carpenter AE, Jones TR, Lamprecht MR, et al. CellProfiler: image analysis software for identifying and quantifying cell phenotypes. Genome Biol 2006;7

Dao D, Fraser AN, Hung J, et al. CellProfiler Analyst: interactive data exploration, analysis and classification of large biological image sets. Bioinformatics 2016;32:3210–12.

Fatkin D, MacRae C, Sasaki T, et al. Missense mutations in the rod domain of the lamin A/C gene as causes of dilated cardiomyopathy and conduction-system disease. N Engl J Med 1999;341:1715–24.

Gerdes AM, Capasso JM. Structural remodeling and mechanical dysfunction of cardiac myocytes in heart failure. J Mol Cell Cardiol 1995;27:849–56.

Hocker JD, et al. 2021. Cardiac cell type–specific gene regulatory programs and disease risk association. Sci Adv 2021;7. 10.1126/sciadv.abf1444

Kanemaru K, Kobayashi Y, Bhaduri A et al. Spatially resolved multiomics of human cardiac niches. Nature 2023;619:801–10.

Koenig AL, et al. 2022. Single-cell transcriptomics reveals cell-type-specific diversification in human heart failure. Nat Cardiovasc Res 1:263–280. 10.1038/s44161-022-00028-6

Litvinukova M, Talavera-López C, Maatz H, et al. Cells of the adult human heart. Nature 2020;588:466–72.

Mahbod A, Schaefer G, Wang C, et al. NuInsSeg: A fully annotated dataset for nuclei instance segmentation in H&E-stained histological images. Sci Data 2024;11:295.

McCain ML, Sheehy SP, Grosberg A, et al. Recapitulating maladaptive, multiscale remodeling of failing myocardium on a chip. Proc Natl Acad Sci U S A 2013;110:9770–75.

Melleby AO, Romaine A, Aronsen JM, et al. A novel method for high precision aortic constriction that allows for generation of specific cardiac phenotypes in mice. Cardiovasc Res 2018;114:1680–90.

Mollova M, Bersell K, Walsh S, et al. Cardiomyocyte proliferation contributes to heart growth in young humans. Proc Natl Acad Sci U S A 2013;110:1446–51.

Muchir A, Shan J, Bonne G, et al. Activation of MAPK pathways links LMNA mutations to cardiomyopathy in Emery-Dreifuss muscular dystrophy. J Clin Invest 2007;117:1282–93.

Otsuji TG, Minami I, Kurose Y, et al. Progressive maturation in contracting cardiomyocytes derived from human embryonic stem cells: qualitative effects on electrophysiological responses to drugs. Stem Cell Res 2010;4:201–13.

Palmer JA, Rosenthal N, Teichmann SA, et al. Revisiting cardiac biology in the era of single cell and spatial omics. Circ Res 2024;134:1681–702.

Skogestad J, et al. Disruption of Phosphodiesterase 3A Binding to SERCA2 Increases SERCA2 Activity and Reduces Mortality in Mice With Chronic Heart Failure. Circulation 2023;147:1221–1236. 10.1161/CIRCULATIONAHA.121.054168

Stringer C, Wang T, Michaelos M, et al. Cellpose: a generalist algorithm for cellular segmentation. Nat Methods 2021;18:100–06.

[dataset] 10x Genomics, 2022, 5k Adult Mouse Heart Nuclei Isolated with Chromium Nuclei Isolation Kit, 10x Genomics https://www.10xgenomics.com/datasets/5k-adult-mouse-heart-nuclei-isolated-with-chromium-nuclei-isolation-kit-3-1-standard (15 May 2026, date last accessed).

[dataset] HuBMAP Consortium, 2021, Azimuth: Human Heart Reference Atlas, HuBMAP/Azimuth Portal, https://azimuth.hubmapconsortium.org/references/#Human%20-%20Heart (8 May 2026, date last accessed).

[dataset] IPA Team, 2023, NuInsSeg: A Fully Annotated Dataset for Nuclei Instance Segmentation in H&E-Stained Histological Images, Kaggle, https://www.kaggle.com/datasets/ipateam/nuinsseg (5 May 2026, date last accessed).

[dataset] Kancherla SK, 2026, CardioSeg, GitHub, https://github.com/SrijanKancherla/CardioSeg (14 May 2026, date last accessed).

